# Mechanism of branched actin assembly at microtubule tips downstream of Adenomatous Polyposis Coli (APC) protein

**DOI:** 10.64898/2026.08.01.742206

**Authors:** Xingyuan Fang, Nadia Efimova, Tatyana M. Svitkina

## Abstract

During cell migration, branched actin filaments nucleated by the Arp2/3 complex induce leading edge protrusions, whereas directionality of cell migration is controlled by microtubules. We showed previously that Adenomatous Polyposis Coli (APC) initiates branched actin assembly at microtubule tips, which can explain how microtubules control directional protrusion. Here, we investigate a link from APC to Arp2/3 complex activity. We show that protrusion-generating activity of APC resides in its N-terminal region containing the Armadillo Repeat Domain (ARD). Furthermore, Asef1/ARHGEF4, a Cdc42 GEF known to be activated by the ARD of APC, as well as Cdc42 itself and its effector N-WASP, an Arp2/3 complex activator, are all required for the assembly of branched actin filaments in neuronal growth cones and neurite outgrowth. As downregulation of Asef1, Cdc42 or N-WASP produces phenotypes similar to APC knockdown, we propose that microtubules regulate directional cell migration and neuron navigation by inducing local membrane protrusion through the APC – Asef1 – Cdc42 – N-WASP – Arp2/3 pathway. These findings bridge a significant gap in our knowledge of cell migration.

## INTRODUCTION

Directional cell migration is a fundamental process in the development and maintenance of multicellular organisms but also plays adverse roles in disease^1, 2^. Precise navigation is especially critical for neurons, which extend neurites over long distances to reach proper targets, whereas navigation failure can lead to neurological disorders^3, 4^.

Actin cytoskeleton with its ability to generate both pushing and pulling forces is the major driver of cell migration^5^. The pulling force is generated by myosin II motors tugging on actin filaments and is used to stabilize cell-substrate adhesions at the cell front and retract the trailing end of migrating cells. The pushing force is generated by polymerizing actin filaments at the leading edge to produce membrane protrusions^6^. The two main types of leading edge protrusions – finger-like filopodia and sheet-like lamellipodia – are driven by parallel bundles of linear actin filaments and networks of branched actin filaments, respectively^5^. Numerous growing actin filament barbed (fast) ends in the branched network allow lamellipodia to generate large force and thus serve as the major protrusive device at the cell front^7, 8^. Most of the barbed ends in lamellipodia are produced by Arp2/3 complex-dependent branched actin nucleation^9–11^.

In neurons, both filopodia and lamellipodia are enriched at the peripheral region of the growth cone, a highly mobile structure at the tip of growing neurites that drives neurite outgrowth and responds to guidance cues in the environment in order to navigate the growing neurites to proper synaptic targets^4, 12^. Many guidance cues induce signaling pathways that eventually lead to activation of the Arp2/3 complex by nucleation promoting factors (NPFs)^13, 14^. The main NPFs responsible for leading edge protrusion are WAVE2 and N-WASP, which in turn are activated by the Rho family GTPases, Rac1 and Cdc42, respectively. The Rac1 ‒ WAVE2 ‒ Arp2/3 complex pathway is the main mechanism to induce lamellipodia for directional cell migration^15, 16^. The Cdc42 ‒ N-WASP ‒ Arp2/3 complex pathway is mostly involved in endocytosis^17^ and invadopodia formation^18^ but can also be implicated in leading edge protrusion^19^, where it is thought to contribute to filopodia formation^20^.

Although regulation of the actin cytoskeleton alone appears sufficient for directional migration of small cells, such as neutrophils^21^ and fish epidermal keratocytes^22^, directional cell migration and front-rear polarity of larger cells, such as fibroblasts^23, 24^ and neurons^25^, depend on microtubule functions.

The best studied mechanisms by which microtubules can control directional cell migration are those involved in regulation of RhoA small GTPase and in focal adhesion disassembly. Microtubules can bind and inactivate GEF-H1, a guanine nucleotide exchange factor (GEF) that activates RhoA^26^. Upon microtubule depolymerization, GEF-H1 is released and becomes active resulting in activation of RhoA, which leads to an increase in actomyosin contractility. The ability of microtubules to target focal adhesions and promote their disassembly^27^ relies on the same GEF-H1 ‒ RhoA ‒ ROCK pathway^28^, although motor-driven delivery of focal adhesion-disassembling cargo along microtubules can also contribute^29^. However, both of these mechanisms contribute to directional migration of nonneuronal cells by promoting deadhesion and retraction of the cell rear and are not obviously implicated in leading edge protrusion. In neurons, both large-scale contraction and deadhesion appear less important, because growth cones have only small punctate adhesions^30^, which quickly turn over without an obvious need for an input from microtubules^31^. Also, neurons do not translocate their body during neurite development and thus do not need large contractile forces.

The question of whether and how microtubules regulate leading edge protrusion remains largely open. Phenomenologically, it was observed that a subset of “pioneer” microtubules approaching the leading edge predict the turning direction of the growth cone^32, 33^, suggesting the existence of such regulatory mechanisms. It is commonly assumed that pioneer microtubules serve as tracks for vesicular transport, which delivers “motility factors”, such as the membrane itself or vesicle-bound signaling molecules^34^. Alternatively, motility factors can reach the leading edge by hitchhiking on the complex of the microtubule plus-end tracking proteins (+TIPs)^35, 36^.

We recently discovered that one of the +TIPs, Adenomatous Polyposis Coli (APC), plays a key role in linking microtubule dynamics to the leading edge protrusion in neuronal growth cones^36^. Using platinum replica electron microscopy (PREM) that is uniquely capable of visualizing fine architecture of the cytoskeleton, we found that a subset of microtubules is associated with a branched actin network at their APC-positive plus ends and that APC knockdown results in nearly complete elimination of branched actin filaments in neuronal growth cones. By live cell imaging we also observed that an encounter of an APC-positive microtubule tip with the plasma membrane was quickly followed by a local actin-rich protrusion^36^. This discovery provides a straightforward answer to the question of how pioneer microtubules can induce membrane protrusion. The next question is how APC mediates this microtubule-branched actin crosstalk. The current work is aimed at answering this question.

APC is best known as a tumor suppressor that functions in the Wnt signaling pathway^37^. Accordingly, APC mutations are linked to various cancer types such as colon and pancreatic cancer^38^. However, APC is also a cytoskeletal protein that has been reported to interact directly with microtubules, actin and intermediate filaments^39–41^. These cytoskeletal functions, rather than Wnt signaling, most likely explain the association of APC mutations with neurological disorders^42–44^.

As a multi-domain protein, APC can be roughly divided into N-terminal, middle and C-terminal regions^45^ (Figure 1A). The unstructured middle region is mainly responsible for APC participation in Wnt signaling through being a part of the β-catenin destruction complex^37^. The C-terminal region of APC (C-APC), which has been predominantly linked to cytoskeletal functions of APC, contains the basic domain that directly binds the microtubule lattice^46^. Downstream of the basic domain, C-APC also has two SxIP sequences that can bind the end-binding (EB) proteins and thus enable APC to act as a +TIP and track the growing ends of microtubules^47^. In addition, two regions within the basic domain are critical for the actin monomer binding and actin filament nucleation *in vitro*^48, 49^. C-APC also contains binding sites for formin mDia1, which helps to elongate the APC-nucleated actin filaments, thus forming long unbranched filaments *in vitro*^49, 50^. Although the actin-nucleating ability of the APC C-terminal regions seems most relevant to the formation of actin-based protrusions, these *in vitro* properties do not explain how APC can initiate the Arp2/3 complex-dependent branched actin filaments at the microtubule tips, as observed in cells^36, 51^.

**Fig. 1.**
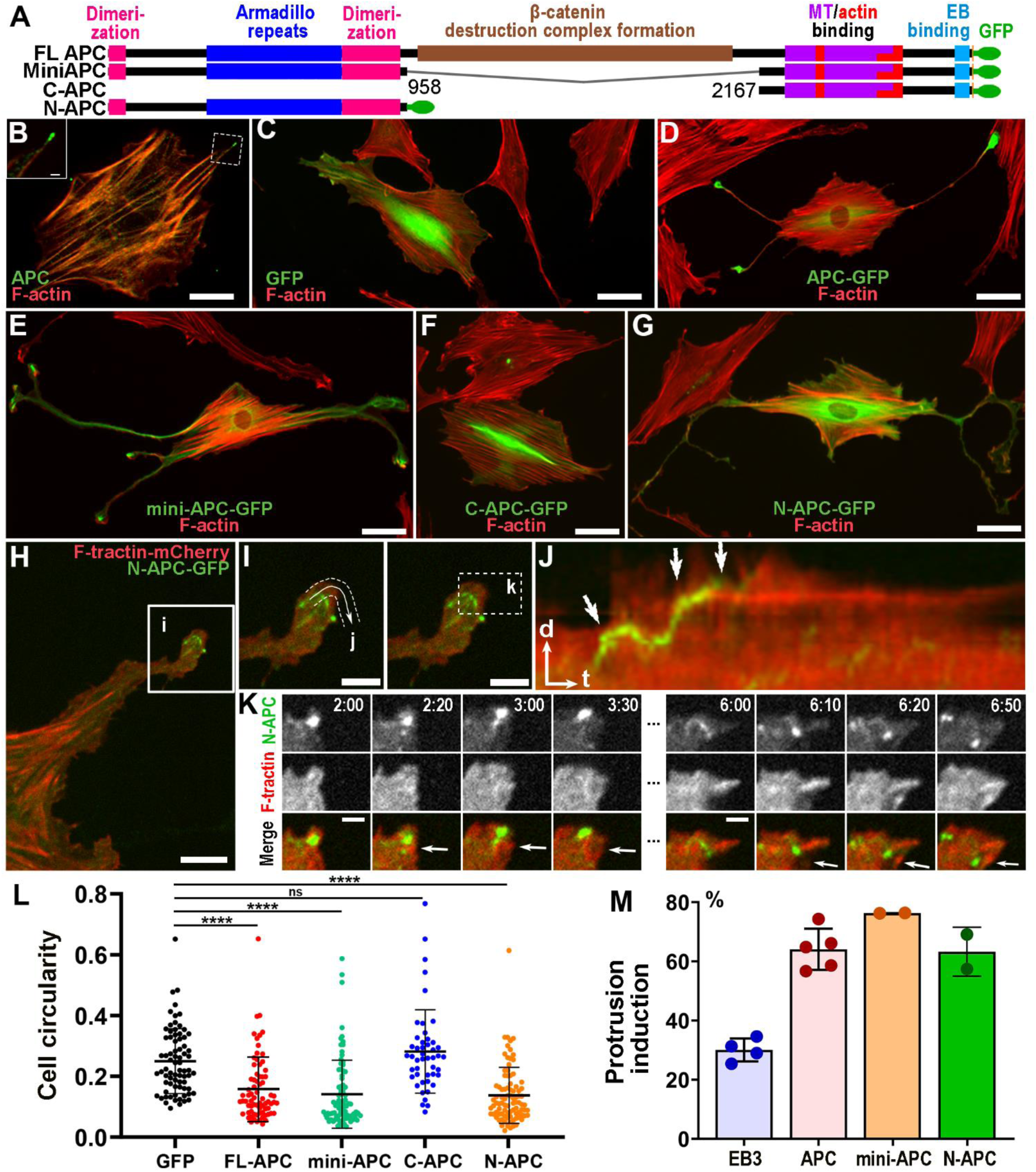
APC induces neurite-like processes in REF52 cells. (A) Diagrams of APC constructs used in the experiments. Amino acid numbers at truncation sites are shown. (B) REF52 cell stained with APC antibody (green) and phalloidin (red). Boxed region is enlarged in inset. Inset: enlarged view of APC cluster at the tip of the process. (C-G) REF52 cells expressing indicated constructs and stained with GFP antibody (green) and phalloidin (red). (H) Process of REF52 cell expressing N-APC-GFP (green) and F-tractin-mCherry (red). (I) Enlarged views of the process tip showing the linear path (j) used to generate the kymograph in J (left) and region of interest (k) for time-lapse frames in K (right). (J) Kymograph of N-APC-GFP (green) and F-tractin-mCherry (red) along the path shown in panel I (left). Encounters of the N-APC punctum with the cell edge (arrows) are followed by F-actin protrusions. (K) Time-lapse frames for the boxed region in panel I (right). Arrows mark the actin-rich protrusion formed after the N-APC-GFP encounters the cell edge. Time is shown in min:sec. (L) Circularity index of cells expressing indicated constructs. Error bars: mean ± SD; ****, p < 0.0001; ns, not significant; Kruskal-Wallis multiple comparisons test with post-hoc Dunn’s test. N = 76 (GFP), 74 (FL-APC), 87 (mini-APC), 48 (C-APC), and 91 (N-APC) cells from three independent experiments. (M) Percentage of EB3-edge or APC-edge encounters followed within ∼30 s by actin-rich protrusions. Data for FL-APC and EB3 are reproduced from our previous study^36^. Error bars: mean ± SD; n = 4 (EB3), 5 (APC) and 2 (mini-APC and N-APC) independent experiments with a total number of 588 APC tracks, 545 EB3 tracks, 131 mini-APC tracks and 122 N-APC tracks. Scale bars: 50 µm (B-G), 10 µm (H), 5 µm(I), and 2 µm (K). Arrows in J: time (t) = 60 s; distance (d) = 2 µm.

As compared with the middle and C-terminal regions of APC, its N-terminal segment (N-APC) appears most promising with regard to branched actin nucleation. Besides several self-association domains^52^, N-APC contains the Armadillo repeat domain (ARD), which is involved in multiple protein-protein interactions^53^. In particular, the ARD binds and activates Asef1^54^, a Cdc42 GEF^55^. Accordingly, we hypothesized that APC activates Arp2/3-dependent branched actin assembly at microtubule tips through the Asef1 ‒ Cdc42 ‒ N-WASP – Arp2/3 complex pathway.

In this study, we tested this model and showed that N-APC is necessary and sufficient for the formation of neurite-like processes in REF52 fibroblasts and that downregulation of Asef1, N-WASP or Cdc42 almost completely eliminated branched actin filaments in growth cones of cultured rat hippocampal neurons, as well as inhibited neurite outgrowth during early neuron development. In conclusion, our study reveals a molecular mechanism downstream of APC that controls the assembly of branched actin filament networks at microtubule tips. These findings bridge the main gap of our knowledge regarding how microtubule can regulate leading edge protrusion.

## RESULTS

### Induction of neurite-like processes by APC overexpression in fibroblasts

To understand the mechanism by which APC initiates branched actin assembly at the microtubule tips, we first aimed to find out which part of APC can produce a gain-of-function phenotype (Fig. 1). For this purpose, we used REF52 cells, which are among a few cell lines that can tolerate ectopic APC expression^56^. REF52 cells express endogenous APC, which is enriched at the tips of cellular processes (Figure 1B, inset), suggesting its role in forming protrusions. Similar localization was reported for other cell types^40, 57, 58^.

Overexpression of GFP-tagged full-length APC in REF52 cells induced long neurite-like processes (Figure 1D). Such processes were rarely seen in non-expressing cells or cells expressing an empty GFP vector (Figure 1C). Using a cell circularity index to quantitatively assess the APC-induced phenotype, we found that the circularity index of cells expressing GFP-APC was significantly lower than that of control cells expressing GFP alone (Figure 1L). This gain-of-function phenotype suggests that full-length APC generates protrusive force for process formation and elongation, likely through Arp2/3 dependent branched actin assembly.

To determine which part of APC is critical for the formation of neurite-like processes, we used APC deletion mutants (Figure 1A). First, to evaluate the possibility that the APC overexpression phenotype results from mis-regulation of β-catenin-dependent signaling, we generated the mini-APC construct by deleting the β-catenin-regulating middle region of APC (Figure 1A). When expressed in REF52 cells, mini-APC, like full-length APC, induced the formation of long neurite-like processes (Figure 1E). The circularity index of cells expressing mini-APC was also significantly lower than that of control cells (Figure 1L). These results suggest that APC induces cell processes using its N-and/or C-terminal domains rather than by modifying β-catenin-dependent transcription. To determine which of the two terminal APC segments is responsible for the gain-of function phenotype, we expressed isolated N-terminal (N-APC) and C-terminal (C-APC) regions of APC in REF52 cells (Figure 1A). The results showed that the expressed C-APC was unable to induce the neurite-like processes in REF52 cells (Figure 1F) or reduce cell circularity (Figure 1L). In contrast, N-APC efficiently induced long processes in cells (Fig. 1G) and reduced cell circularity similar to full-length APC and mini-APC (Figure 1L). These results show that the ability of APC to stimulate neurite-like processes in REF52 cells mainly depends on its N-terminal part, but not on the C-terminal region despite the ability C-APC to facilitate F-actin assembly in vitro^50^ and increase microtubule stability in cells^59^.

### Induction of local actin-rich protrusions by APC-positive microtubules

The ability of N-APC to stimulate neurite-like processes in REF52 cells suggests that it can induce actin assembly at the cell leading front. The expression of fluorescently labeled APC constructs offered an opportunity to correlate APC dynamics with actin-based protrusions in cells. We previously showed that when full-length APC riding on a growing microtubule tip hit the cell edge, this encounter was quickly followed by a transient actin-rich protrusion in 64.1% of cases, whereas such events were much less common (30.1%) when EB3-tipped microtubules were monitored^36^.

Here, we used the same approach to determine whether mini-APC and N-APC can also induce protrusions. To this end, we co-expressed N-APC or mini-APC with mCherry-F-tractin, a marker of F-actin, in REF52 cells (Figure 1H-K). We found that both N-APC and mini-APC, similar to full-length APC, formed dynamic accumulations at the tips of cellular processes (Figure 1H; Supplemental Figure S1A). Within these accumulations, individual APC clusters, likely associated with microtubule tips, made relatively short runs toward the cell edge, or roamed around within a small area, or remained nearly stationary (Videos 1 and 2). During their roaming behavior, APC clusters usually remained distant from the cell edge. No obvious correlation between APC dynamics and cell edge protrusion was observed in such cases. The stationary APC clusters were also unsuitable for correlative analysis. Therefore, we focused on the events when an APC mutant punctum hits the cell edge after a detectable run toward the edge. As in the case of full-length APC^36^, the encounters of N-APC or mini-APC with the cell edge were often followed by a local actin-rich protrusion in REF52 cells (Figure 1H-K; Supplemental Figure S1A-E) with the frequencies comparable to that exhibited by full-length APC, namely 61.9% for N-APC and 76.3% for mini-APC (Figure 1M). These results show a strong correlation between the dynamics of N-terminus-containing APC constructs and the formation of actin-rich protrusions at the leading edge, suggesting a causative role of the N-terminal region of APC in this process.

### Endogenous APC is not necessary for induction of neurite-like processes by N-APC

The ability of N-APC-positive microtubule tips to induce actin-rich protrusions is consistent with our hypothesis that the ARD within N-APC triggers branched actin polymerization by activating the Asef1 – Cdc42 – N-WASP pathway. However, besides ARD, the N-APC construct also contains several self-association domains^45^, which have been shown to mediate APC dimerization both *in vitro* and in cells^48, 60^. Thus, an alternative explanation for N-APC-mediated induction of actin-rich protrusions (Figure 1H-K) is that ectopic N-APC dimerizes with endogenous APC, and it is the endogenous full-length APC that induces neurite-like processes, for example, by using its actin-nucleating capability^49, 50^. To address this possibility, we knocked down endogenous APC in REF52 cells using previously validated lentivirus-encoded shRNAs with Cerulean fluorescent protein as a reporter^36^ and then expressed shRNA-resistant versions of either N-APC or C-APC in these cells (Figure 2).

**Fig. 2.**
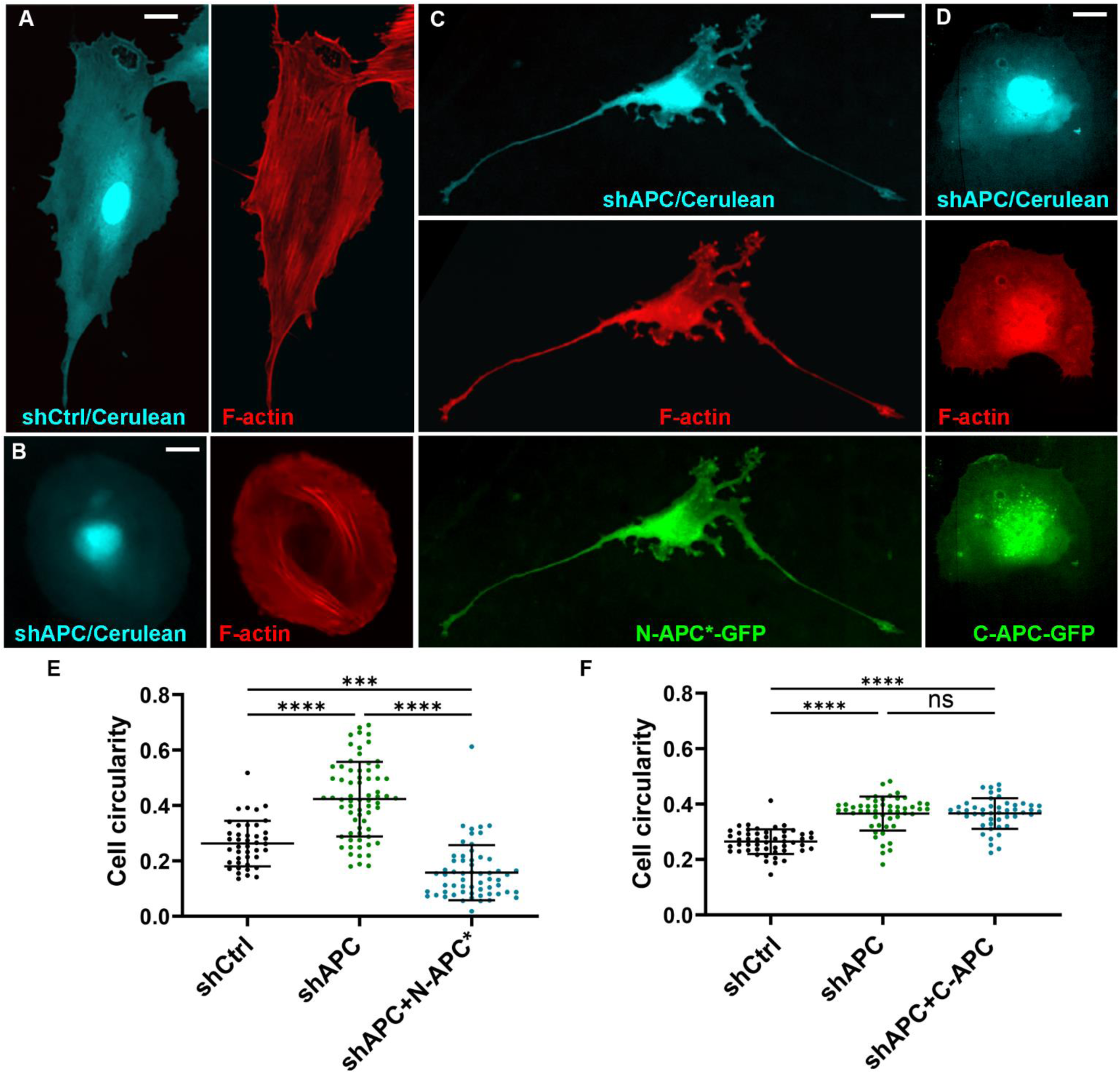
N-APC, but not C-APC, is sufficient to promote extension of neurite-like processes in REF52 cells. (A-D) REF52 cells were transfected with control (shCtrl; A) or APC-targeting (shAPC; B-D) shRNAs together with Cerulean fluorescent protein (cyan) for 4 days and rescued with none (B), or shRNA-resistant N-APC*-GFP (C, green) or C-APC-GFP (D, green), fixed and stained with phalloidin (A-D, red). Scale bars: 20 µm. (E, F) Circularity index of cells expressing indicated shRNAs with or without N-APC*-GFP (E) or C-APC-GFP (F), as indicated. Error bars: mean ± SD; ****, p < 0.0001; ***, p < 0.001; ns, not significant; Kruskal-Wallis multiple comparisons test with post-hoc Dunn’s test. N = 42 (shCtrl), 68 (shAPC), and 59 (shAPC+N-APC*) cells for graph in E and 51 (shCtrl), 51 (shAPC), and 47 (shAPC+C-APC) cells for graph in F from three independent experiments for each graph.

Remarkably, APC depletion in REF52 cells resulted in a loss of virtually all elongated processes and even sharp corners in the cell shape, so that the cells became significantly rounder (Figure 2B) than control cells treated with nontargeting control shRNA against Luciferase (Figure 2A). These observations were validated by quantification of the cell circularity index, which was found to be significantly higher for APC knockdown cells than for control cells (Figure 2E, F). Additionally, F-actin organization in cells became more diffuse with fewer prominent stress fibers.

When expressed in the APC knockdown cells, N-APC was still able to induce neurite-like processes (Figure 2C) suggesting that this effect was independent of endogenous full-length APC but was mediated by activities residing within N-APC itself. On the other hand, the expression of C-APC failed to change the roundish shape of APC knockdown cells or restore F-actin stress fibers (Figure 2D). In fact, C-APC seemed to exacerbate the latter phenotype, making F-actin distribution largely diffuse throughout the cell except for F-actin accumulation in lamellipodia and ruffles. These results strongly suggest that N-APC induces neurite-like processes using its own activities, without the help of endogenous APC.

### Knockdown of Asef1, Cdc42 or N-WASP in rat hippocampal neurons reduces F-actin levels in growth cones

Combining our current data showing that N-APC promotes actin-rich protrusions in REF52 cells (Figures 1 and 2) with our previous findings that APC is required for the assembly of branched actin networks in neuronal growth cones^36^, we propose that N-APC is chiefly responsible for the APC-dependent branched actin assembly. In cells, where full-length APC interacts with microtubule tips using its C-terminal sequences, the N-APC region can trigger branched actin assembly at microtubule tips by activating the Asef1 – Cdc42 – N-WASP – Arp2/3 complex signaling and thereby induce membrane protrusions.

To test this hypothesis, we evaluated roles of Asef1, Cdc42 and N-WASP in neuronal growth cones. We first used immunofluorescence staining of dissociated rat hippocampal neurons cultured for 7 days in vitro (DIV7) to determine localization of these three proteins. We found significant enrichment of Asef1, Cdc42 and N-WASP in actin-rich growth cones (Figure 3A-C), suggesting that these proteins are in the right place to be able to function as the downstream effectors of N-APC.

**Fig. 3.**
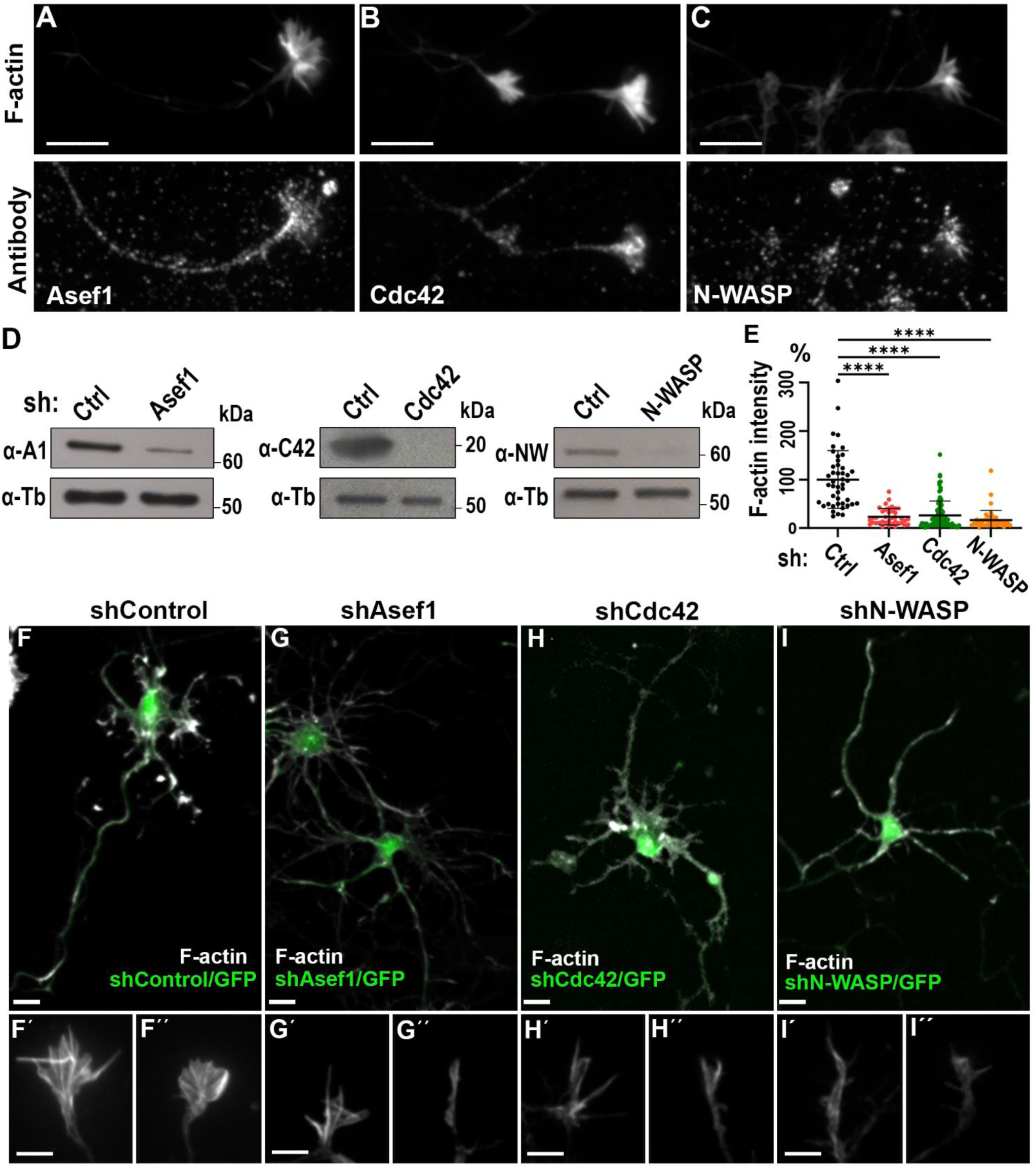
Asef1, Cdc42 and N-WASP localize to growth cones of cultured rat hippocampal neurons and are important for growth cone morphology. (A-C) Fluorescence staining of DIV7 hippocampal neurons with phalloidin (F-actin; top panels) and indicated antibody (bottom panels). Scale bars: 10 µm. (D) Western blots of lysates from REF52 cells transfected with control shRNA (Ctrl), or shRNAs targeting Asef1, Cdc42 or N-WASP and probed with Asef1 (α-A1), Cdc42 (α-C42) or N-WASP (α-NW) antibodies as indicated. Tubulin antibody staining (α-Tb) is used as loading control. (E) Quantification of total fluorescence intensity of phalloidin staining in growth cones of neurons treated with indicated shRNAs at DIV1 and imaged at DIV5. Error bars: mean ± SD; N = 38 (Ctrl), 86 (shAsef1), and 45 (shN-WASP) growth cones from three independent experiments. ****, p < 0.0001; Kruskal-Wallis multiple comparisons test with posthoc Dunn’s test. (F-I) Phalloidin staining of DIV5 neurons treated for 4 days with control (shControl) or Asef1-, Cdc42-, and N-WASP-targeting shRNAs with GFP reporter (green). Bottom panels show representative growth cones in each group at high magnification. Scale bars: 20 µm (top panes) and 5 µm (all bottom panels).

Next, we used shRNA-mediated knockdown of either Asef1, Cdc42 or N-WASP in rat hippocampal neurons to determine the functional contribution of these proteins to morphology and actin organization in neuronal growth cones. For this purpose, we used lentiviruses encoding both the shRNA and GFP as a reporter. We designed two shRNAs for each protein, tested their efficiency by Western blotting in REF52 cells, which are also from rat, and chose one shRNA for each protein that showed better knockdown efficiency across three repeats (Figure 3D).

To knockdown Asef1, Cdc42 and N-WASP in rat hippocampal neurons, we infected neurons with lentiviral shRNAs at DIV1 and fixed them at DIV5. As revealed by phalloidin staining, control neurons had normal fan-shaped growth cones enriched with F-actin (Figure 3F). In contrast, the growth cones in Asef1, Cdc42 and N-WASP knockdown neurons were smaller and displayed a spiky shape most often characterized by a single narrow actin-containing region at the tips of the neurite or sometimes consisting of a few filopodia with undetectable lamellipodia in-between (Figure 3G-I). These altered growth cone shapes suggest deficient ability of knockdown neurons to form lamellipodia, which are responsible for the broad fan-like shape of growth cones, whereas filopodia appeared less affected. Quantitative evaluation of actin-rich regions at the neurite tips, revealed a significant decrease in the average fluorescence intensity of phalloidin staining, as well as in the growth cone area, after knockdown of Asef1, Cdc42 or N-WASP, as compared with the control group (Figure 3E). Similar reductions in growth cone F-actin intensity and size were also observed for the other, lower efficiency shRNAs targeting Asef1 and N-WASP, whereas the second shRNA for Cdc42 was inefficient by Western blotting and was not tested in neurons.

Together, the results of knockdown experiments demonstrated that Asef1, Cdc42 and N-WASP play key roles for the assembly of a normal levels of F-actin in growth cones and, specifically, in the formation of growth cone lamellipodia, which are rich in branched actin filaments in contrast to filopodia. Importantly, the similar phenotypes resulting from knockdown of each of these proteins suggest that Asef1, Cdc42 and N-WASP function is the same pathway.

### Asef1, Cdc42 or N-WASP knockdown leads to a loss of branched actin filaments in neuronal growth cones

Although the preferential loss of growth cone lamellipodia observed by fluorescence microscopy after knockdown of Asef1, Cdc42 or N-WASP in hippocampal neurons suggested that these proteins are specifically important for the assembly of branched actin filaments, resolution of light microscopy is insufficient to provide direct evidence on this point. Therefore, we next used PREM to determine which subsets of actin filament arrays are impaired in Asef1-, Cdc42-or N-WASP-depleted growth cones (Figure 4).

**Fig. 4.**
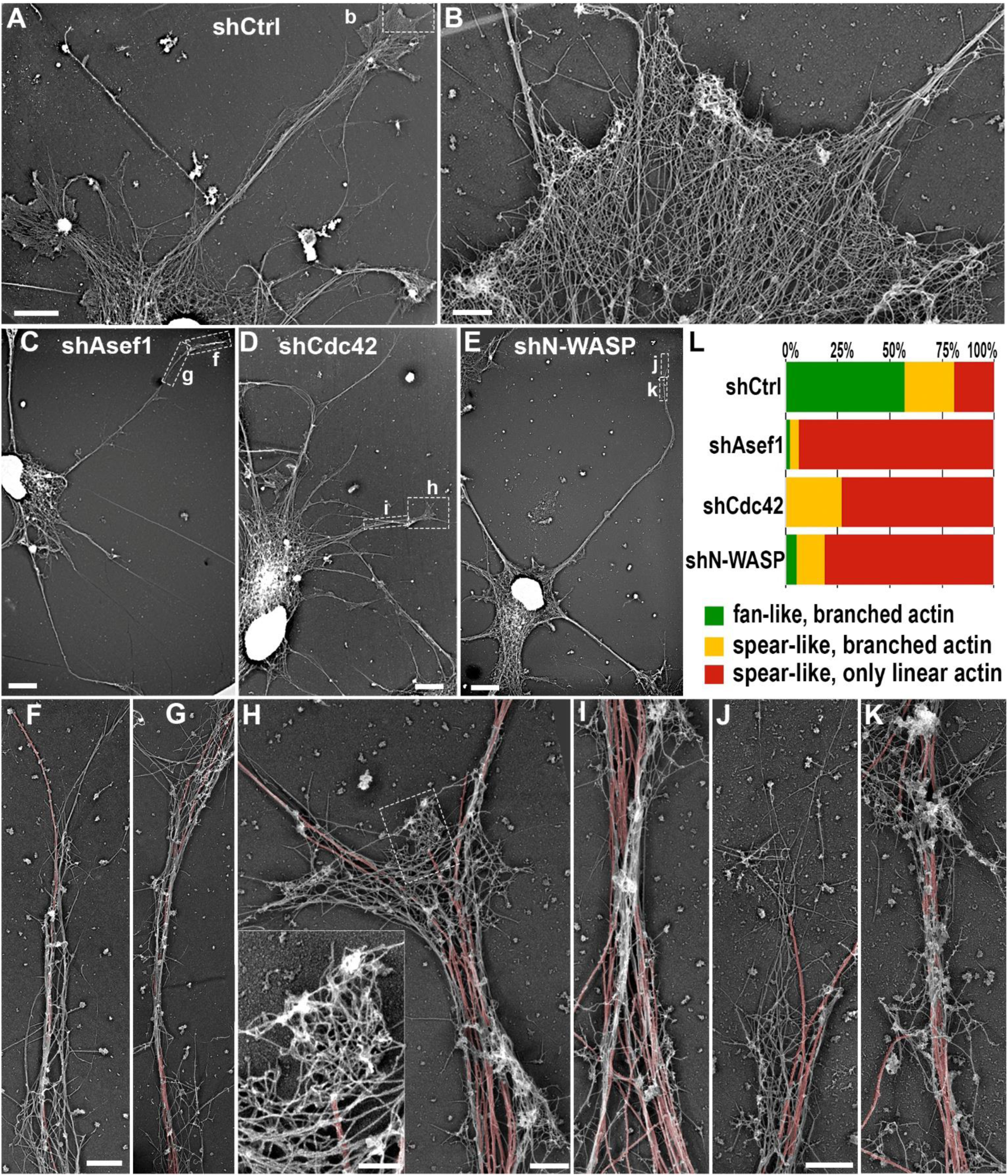
Depletion of Asef1, Cdc42 or N-WASP suppresses branched actin filaments in growth cones without significantly affecting linear actin filaments. (A-K) PREM images of rat hippocampal neurons treated with control shRNA (A, B) or shRNAs targeting Asef1 (C, F, G), Cdc42 (D, H, I) or N-WASP (E, J, K). (A, C, D, E) Overviews showing significant portions of DIV5 neurons treated with indicated shRNAs. Boxes labeled with low-case letters indicate regions enlarged in respective main panels. (B) Enlarged growth cone from control neuron showing dense branched actin. (F, G) Enlarged view of spear-shaped neurite tip from (C) with only linear actin filaments. (H, I) Enlarged view of growth cone-like structure from (D) with mostly linear actin filaments and a small patch of branched actin (box), which is enlarged in the inset. (J, K) Enlarged view of the neurite tip from (E) with only linear actin filaments. Microtubules in F-K are highlighted in red. (L) Percentages of structural categories observed at neurite tips of shRNA-treated neurons. N = 21 (shCtrl), 48 (shAsef1), 41 (shCdc42), and 37 (shN-WASP) neurite tips from 7, 9, 8, and 9 cells, respectively, from three independent experiments. Scale bars: 5 µm (A, C, D, E), 0.5 µm (B, F-K) and 100 nm (H, inset).

Control growth cones of DIV5 neurons, as expected, typically had a fan-like shape and contained extensive networks of branched actin filaments, as well as long unbranched actin filaments, either in the form of filopodial bundles or as a part of lamellipodia (Figure 4A, B).

After depletion of Asef1, Cdc42 or N-WASP, the growth cones almost invariably exhibited either a spear-like shape or a fork-like configuration, consistent with the fluorescence microscopy observations. Strikingly, the neurite tips in knockdown neurons were dramatically deprived of branched actin networks and mostly contained linear actin filaments interspersed with microtubules (Figure 4C-K). When detected at the neurite tips in knockdown neurons, the branched actin networks were typically present in the form of small patches, which were often associated with microtubule tips (Figure 4H, inset). Given the overall severity of branched actin depletion phenotypes, we speculate that remaining branched actin patches reflect incomplete knockdown of the target protein.

To quantify the PREM phenotypes after knockdown of Asef1, Cdc42 or N-WASP, we categorized the neurite tips into three types based on their shape and actin filament architecture: 1) fan-shaped growth cones with extensive branched actin network (e.g.: Figure 4B); 2) spear-shaped or fork-shaped growth cones with small patches of branched actin filaments (e.g.: Figure 4H); and 3) growth cones with only linear actin filaments (e.g.: Figure 4F, J). In control neurons, 57.1% growth cones belonged to the first category and only 19% growth cones contained only linear actin filaments (category 3). These category 3 neurite tips were probably not in a state of active protrusion. In contrast, after knockdown of either Asef1, Cdc42 or N-WASP, neurite tips with only linear actin filaments were predominant and comprised 93.7%, 73.2% and 81.1%, respectively (Figure 4L).

The severe loss of branched actin networks after Asef1, Cdc42 or N-WASP depletion demonstrates that these three proteins are crucial for the assembly of branched actin networks in growth cones. Since these knockdown phenotypes closely mimic the effects of APC depletion^36^, Asef1, Cdc42 or N-WASP likely act downstream of APC to induce branched actin filament assembly in growth cones.

### Acute inhibition of Cdc42 or N-WASP inhibits neurite outgrowth

Given that branched actin networks form the core mechanism underlying leading edge protrusion in migrating cells, including the outgrowth of neuronal processes driven by growth cone motility^61–63^, it can be expected that Asef1, Cdc42 or N-WASP play important roles in neurite outgrowth. Since shRNA-mediated knockdown takes relatively long time, neurites can achieve significant lengths by the time the depletion of a target protein produces an effect, which makes detection of the neurite outgrowth phenotype difficult. Therefore, we utilized a pharmacological approach to acutely inhibit protein functions at an early stage of neuronal development (DIV1). Chemical inhibitors ML141 and wiskostatin are available for Cdc42 and N-WASP, respectively^64, 65^.

The use of pharmacological inhibitors also offers an alternative approach, besides protein knockdown, to test roles of these proteins in growth cone structure. Therefore, we first characterized the effect of Cdc42 or N-WASP inhibition on the morphology and actin organization in neuronal growth cones. Neurons were cultured for 4-6 hours after plating and then treated with either 20 μM ML141 or 6 μM wiskostatin for 18-20 hours to inhibit Cdc42 or N-WASP, respectively, or with 0.1% DMSO as control (Figure 5). Control neurons at this stage (DIV1) usually developed a small number of neurites tipped with actin-rich fan-shaped growth cones but also retained broad actin-rich lamellipodia associated with the soma, which are a dominant feature of very young neurons (Figure 5A; Supplemental Figure S2A).

**Fig. 5.**
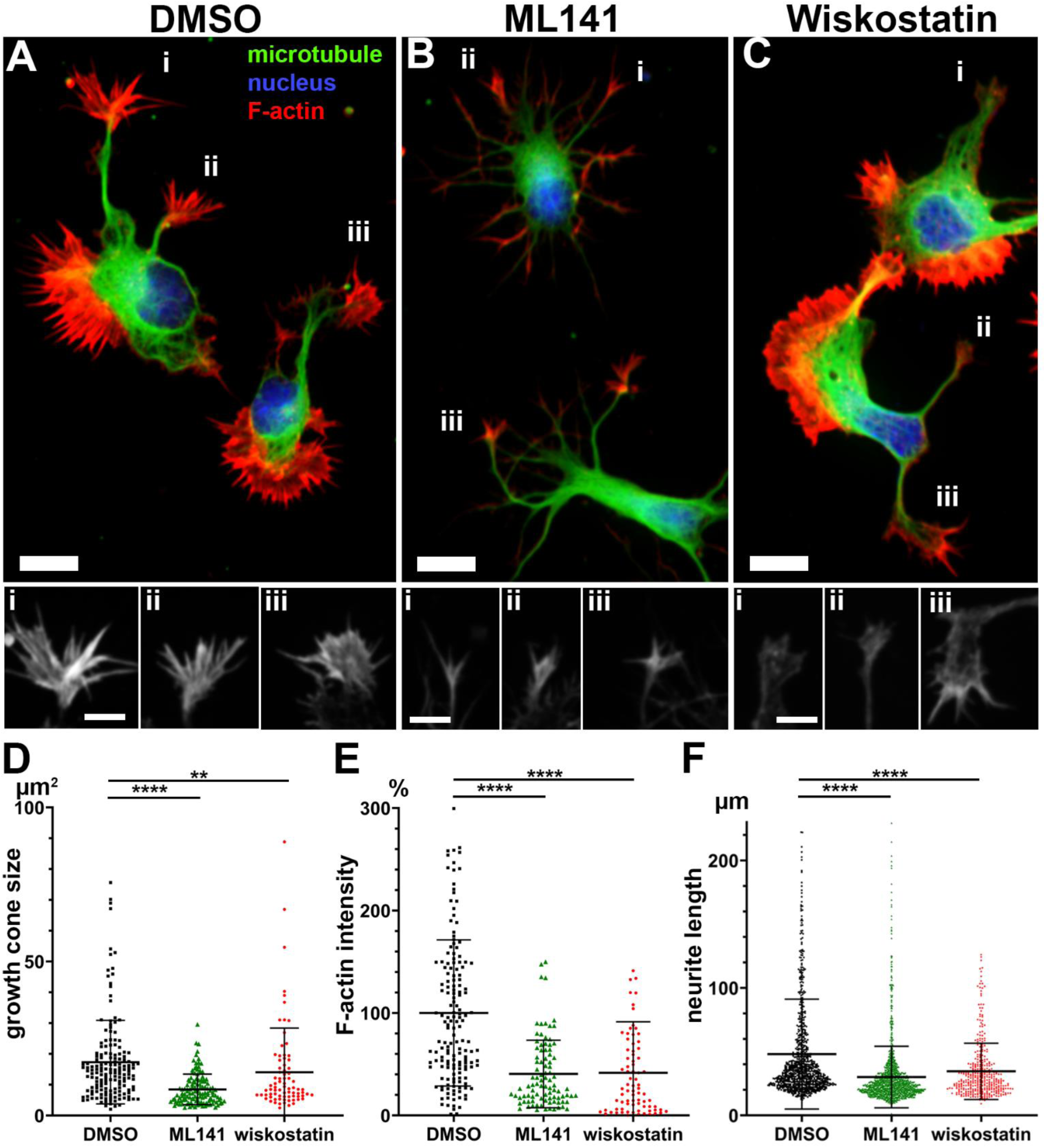
Pharmacological inhibition of Cdc42 and N-WASP reduces growth cone size, F-actin intensity in growth cones and neurite lengths in rat hippocampal neurons. (A-C) Staining of DIV2 neurons with phalloidin (F-actin, red), β3 tubulin antibody (green) and DAPI (blue) after treatment with 0.1% DMSO (A), 20 µM ML141 (Cdc42 inhibitor) or 6 µM wiskostatin (N-WASP inhibitor) for 18-20 hrs. Individual phalloidin-stained growth cones (i, ii, iii) from each panel are shown in bottom panels in greyscale. Phalloidin intensity levels are adjusted identically for different conditions. Scale bars: 10 µm (main panels and 5 µm (lower panels). (D-F) Quantification of growth cone size (D), average fluorescence intensity of phalloidin staining (E) and neurite length (F) in after indicated treatments. Error bars: mean ± SD; ****, p < 0.0001; **, p < 0.01; Kruskal-Wallis multiple comparisons test with posthoc Dunn’s test. N = 168 (DMSO), 133 (ML141), and 85 (wiskostatin) neurite tips for graphs in D and E from three independent experiments; N = 900 (DMSO), 1456 (ML141), and 392 (wiskostatin) neurite tips for graph F from three independent experiments.

Compared with control neurons (Figure 5A), ML141-treated neurons lacked soma-associated lamellipodia, but contained a larger number of neurites (Figure 5B, Supplemental Figure S2B). Phalloidin staining revealed the presence of F-actin at the neurite tips in ML141-treated neurons suggesting the presence of growth cone-like structures. However, compared with control neurons, ML141-treated growth cones had a spiky shape and were smaller with lower intensity of phalloidin staining (Figure 5B, D, E), thus mimicking the phenotype of Cdc42 knockdown neurons (Figure 3H).

Neurons treated with wiskostatin had an overall shape similar to control neurons. They exhibited a small number of neurites with growth cones and broad soma-associated lamellipodia (Figure 5C, Supplemental Figure S2C). Whereas these lamellipodia were indistinguishable from analogous lamellipodia in control neurons, the growth cones at the tips of extended neurites had a smaller size and lower F-actin contents, as compared with growth cones in control neurons (Figure 5C, D, E), although they appeared less spiky than after Cdc42 inhibition and exhibited some broadening relative to the neurite shaft. Thus, N-WASP inhibition had a similar effect on growth cones at the tips of neurites as N-WASP knockdown (Figure 3I), but wiskostatin failed to affect the soma-associated lamellipodia. Since such lamellipodia are characteristic for young neurons at DIV1 but no longer exist at DIV5, the effects of pharmacological and genetic downregulation of N-WASP on growth cones specifically can be considered comparable. We assume that soma-associated lamellipodia, in contrast to growth cones, more critically depend on the Rac1 – WAVE – Arp2/3 complex pathway than on the Cdc42 – N-WASP – Arp2/3 complex pathway.

To take advantage of the ability of drugs to acutely inhibit target proteins, we also evaluated the degree of neurite outgrowth in ML141-and wiskostatin-treated neurons in comparison with control, DMSO-treated neurons. Quantification of the neurite lengths in all three groups demonstrated that inhibition of either Cdc42 or N-WASP inhibited neurite outgrowth (Figure 5F), suggesting that Cdc42 and N-WASP activities have a strong impact not only on the growth cone morphology but also on the growth cone functions in driving neurite outgrowth.

### Inhibition of Cdc42 or N-WASP leads to a loss of branched actin networks in neuronal growth cones

Given the similarity between the knockdown and drug-induced phenotypes, as seen by fluorescence microscopy, we hypothesized that branched actin filament networks could be also lost in growth cones after drug treatments, which would explain the impaired ability of growth cone to extend neurites. To test this hypothesis, we used PREM to visualize the actin cytoskeleton architecture of growth cones after treatment with either ML141 or wiskostatin (Figure 6).

**Fig. 6.**
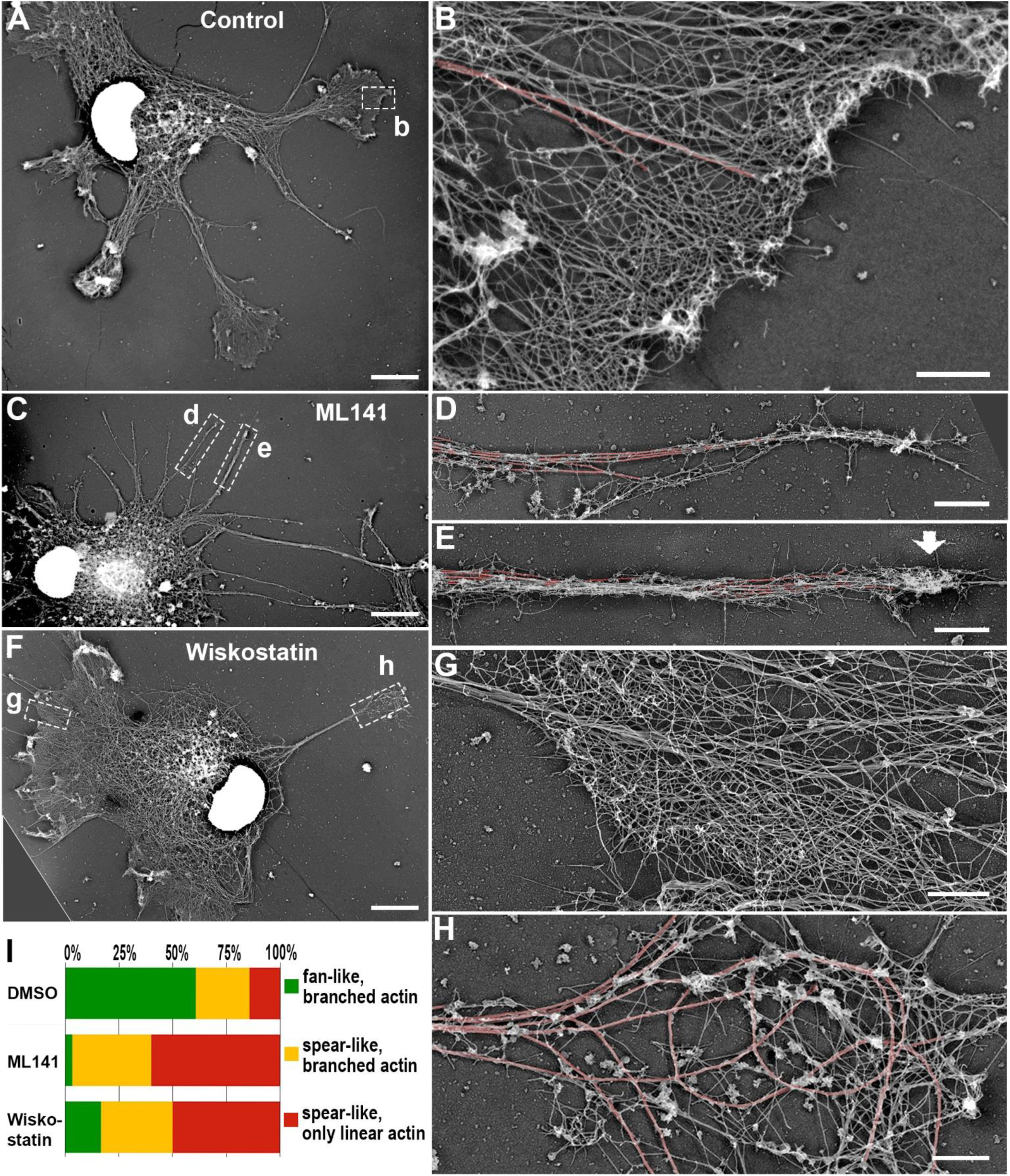
Pharmacological inhibition of Cdc42 and N-WASP suppresses branched actin filaments in growth cones without significantly affecting linear actin filaments. (A-H) PREM images of rat hippocampal neurons treated with 0.1% DMSO (A, B), 20 µM ML141 (C-E) or 6 µM wiskostatin F-H). (A, C, F) Overview images of significant portions of DIV2 neurons treated with indicated drugs. Boxes labeled with low-case letters indicate regions enlarged in respective main panels. (B) Enlarged growth cone lamellipodium of DMSO-treated neuron showing dense branched actin filaments. (D, E) Enlarged spear-shaped neurite tips from ML141-treated neuron containing either only linear actin filaments (D) or linear filaments together with a small patch of branched actin (E, arrow). (G, H) Enlarged views from wiskostatin-treated neuron of soma-associated lamellipodium with dense branched actin filaments (G) and neurite tip with sparse linear actin filaments (H). Microtubules in B, D, E, G, and H are highlighted in red. (I) Percentages of structural categories observed at neurite tips of control or drug-treated neurons. N = 76 (DMSO), 56 (ML141), and 140 (wiskostatin) neurite tips from three independent experiments. Scale bars: 5 µm (A, C, F), 1 µm (B, G) and 2 µm (D, E, H).

Similar to growth cones of control DIV5 neurons (Figure 4A), growth cones of control DMSO-treated DIV1 neurons, had a fan-like shape and contained extensive networks of branched actin filaments (Figure 6A, B). In contrast, the tips of multiple neurites in ML141-treated neurons mostly contained linear actin filaments intermixed with microtubules (Figure 6C-E) with occasional presence of small patches of branched actin filaments (Figure 6E). On the other hand, fan-shaped growth cones with significant amount of branched actin filaments were extremely rare after Cdc42 inhibition (Figure 6I). In wiskostatin-treated neurons (Figure 6F-H), the branched actin network in the broad soma-associated lamellipodia appeared largely intact (Figure 6G), whereas the branched actin network in growth cones at the tips of neurites was largely eliminated (Figure 6H). Despite the somewhat broader shape of wiskostatin-treated growth cones, the growth cone interior was mostly occupied by curved microtubules with only sparse linear actin filaments, while actin filaments were largely restricted to the growth cone periphery and were mostly linear. Using the same categorization of growth cone structure, as described for figure 4, we quantified the effects of drug treatments and found that 60.7% growth cones in control neurons were fan-shaped with branched actin networks, and only 14.8% growth cones were spear-shaped with only linear actin filaments. In contrast, after treatment with either ML141 or wiskostatin, the fraction of fan-shaped growth cones with branched actin filaments decreased dramatically to 3.3% and 16.7%, respectively (Figure 6I). The rest of these populations comprised aberrant growth cones with either linear filaments only or linear filaments together with small patches of branched actin networks. These quantitative results confirm a significant loss of branched actin in growth cones after inhibition of Cdc42 or N-WASP, consistent with the result of respective RNAi treatments. The slightly less dramatic effects of drugs on growth cones, as compared to those resulting from shRNA treatment, could be explained by relatively low concentrations of ML141 and wiskostatin we used in pharmacological experiments in order to keep neurons relatively healthy and minimize off-target effects.

## DISCUSSION

Building on our previous discovery that APC is responsible for branched actin formation at the microtubule tips, as well as throughout the neuronal growth cone^36^, in this study we investigated how APC activity is linked to branched actin polymerization. We show that (1) APC positively regulates protrusions not only in neurons but also in fibroblasts; (2) APC’s protrusion-generating activity resides in the ARD-containing N-terminal region of APC rather than in its microtubule-binding and actin-nucleating C-terminal region; and (3) Asef1, Cdc42 and N-WASP, similar to APC, are necessary for the formation of branched actin filament networks in growth cones and play important roles in the outgrowth of neuronal processes. Based on these findings, we propose a straightforward model of how microtubules navigate both neurite outgrowth and migration of non-neuronal cells (Fig. 7). This model includes the following events: (1) APC tracks growing microtubule tips using the EB-binding and microtubule lattice-interacting sequences in its C-terminal region; (2) when the APC-tipped dynamic microtubule approaches the membrane, it encounters its downstream effectors there; (3) the ARD within the N-terminal region of APC binds and activates the Cdc42 GEF Asef1; (4) this event triggers the Asef1 – Cdc42 – N-WASP – Arp2/3 complex pathway, which leads to local assembly of branched actin filaments; and (5) branched actin filaments push onto the membrane at the site of the microtubule-membrane encounter, thus causing a protrusion at the site pointed out by the microtubule (Fig. 7).

**Fig. 7.**
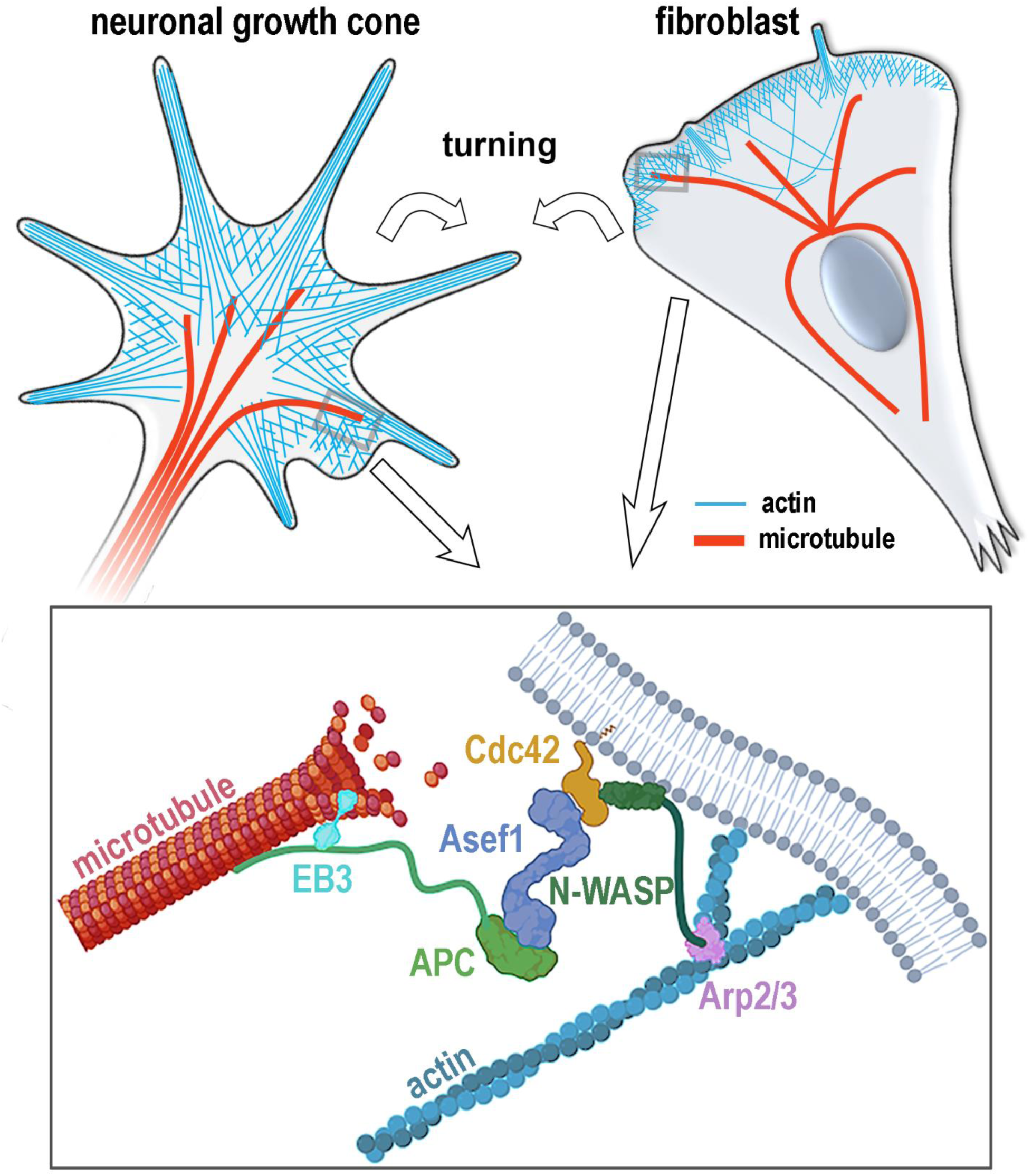
Diagram illustrating the proposed mechanism by which APC can initiate turning of a neuronal growth cone (upper left) or a migratory cell (upper right) by triggering formation branched actin filaments at the microtubule tip through the APC – Asef1 – Cdc42 – N-WASP – Arp2/3 complex pathway (bottom).

Although this model can fit the previously known concepts of the actin-microtubule crosstalk, such microtubule-dependent regulation of small GTPases and delivery of “motility” factors to the cell front^39^, the conceptual advance provided by our findings is in the molecular details of the underlying mechanism.

### APC promotes protrusions in non-neuronal cells

We chose REF52 fibroblasts as a model system to define roles of APC fragments, because overexpression of full-length APC in these cells induces a striking phenotype – formation of long neurite-like processes. These processes are not filopodia, as they are much longer, contain microtubules and are often associated with a dynamic growth cone-like tip, which emits small transient actin-rich protrusions. Besides providing an excellent readout for the APC gain-of-function phenotype, these findings also show that non-neuronal REF52 cells have all necessary machinery to stimulate protrusions downstream of APC when APC is provided in excess. Moreover, this APC-dependent machinery is also endogenously functional in wild-type cells, because knockdown of APC largely eliminates the elongated cellular processes in wild type REF52 cells. Together with our previous observations that branched actin networks can be associated with microtubule tips not only in neurons but also in other cell types^36, 51^, these data suggest that APC is involved in microtubule-dependent protrusion formation not only in neurons but much more broadly.

### N-APC is responsible for the gain-of-function phenotype

Using overexpression of APC deletion mutants in REF52 cells, we show that neither the middle domain of APC, which is involved in β-catenin regulation downstream of Wnt signaling^37^, nor the C-APC fragment that contains the main cytoskeleton-binding sequences^45^, is necessary for the formation of neurite-like processes, whereas the N-APC region containing the ARD, a protein-protein interaction module, is necessary and sufficient for this phenotype. This conclusion is further validated by the ability of N-APC to produce the gain-of-function phenotype in the absence of endogenous APC, thus ruling out a requirement for this effect of C-APC, which could be supplied by endogenous APC in the overexpression experiments.

The inability of C-APC to induce protrusions either after overexpression or in knockdown-rescue experiments may seem surprising in the light of the *in vitro* ability of the basic domain within C-APC to nucleate actin filaments by itself and also cooperate with mDia1 to elongate these unbranched actin filaments^49, 50^. However, it has been shown that APC binding either to EB1 or to the microtubule lattice via its C-terminal sequences conflicts with APC binding to actin monomers^66^, suggesting that when APC is bound to microtubules, actin nucleation by C-APC is blocked. There data make it less surprising that C-APC fails to produce a gain-of-function phenotype in REF52 cells. Potentially, the C-APC-dependent actin nucleation plays other roles in cells than in leading edge protrusion, for example, in focal adhesion regulation^48^.

By expressing deregulated N-APC in APC knockdown cells, we expected to find broad lamellipodia all over the cell perimeter due to predicted inability of N-APC to track microtubule plus ends and thus be targeted by microtubules to specific membrane sites. However, contrary to our expectations, N-APC induced long narrow processes similar to those induced by full length APC, although less well defined. These data suggest that N-APC is still able to be locally concentrated and retained at the cell periphery for a significant amount of time to induce formation of cell processes. One possible explanation for this phenomenon is that N-APC is still targeted by microtubules but rather than tracking their plus ends, it travels along microtubules^67^ as a cargo carried by kinesins^68, 69^. As the ARD of APC can also bind the membrane-associated protein Amer1^70^, this interaction can prolong APC localization at the plasma membrane. Thus, the constant delivery of N-APC by microtubules and its local stabilization at the plasma membrane can sustain persistent protrusion and result in the formation of long processes.

### Downstream effectors of N-APC

Among multiple ARD interaction partners, we chose to test the most promising Asef1 ‒ Cdc42 ‒ N-WASP pathway, because it leads to activation of the Arp2/3 complex, the only protein known to generate branched actin filament networks that we have observed at microtubule tips^36^. Our results show that knockdown of either Asef1, Cdc42 or N-WASP, or pharmacological inhibition of Cdc42 or N-WASP, in hippocampal neurons virtually eliminates branched actin filaments in growth cones, thus closely mimicking the phenotype of APC knockdown^36^. The similarity of the downregulation phenotypes among all four proteins strongly suggests that all of them function in the same pathway and that this pathway is responsible for the formation of branched actin filaments in neuronal growth cones.

Although initially reported to be a Rac-activating GEF, Asef1 and its paralog Asef2 have been subsequently demonstrated to be exclusively Cdc42 activators^54, 55^. Asef1 forms an autoinhibited conformation that can be disrupted by the ARD of APC, thus allowing Asef1 to activate Cdc42^71^. Previous studies have shown that constitutively active Asef1 that lacks an autoinhibitory domain, or full length Asef1 when it is coexpressed with the ARD of APC, induced lamellipodia in non-neuronal cells^54^, suggesting that this phenotype most likely involves the roles of Cdc42 and N-WASP, as in our study.

Cdc42 was initially thought to function mostly in filopodia formation^72^, but now this small GTPase is considered to be a master regulator of cell polarity in different contexts, including cell migration^57, 73^ and neuronal morphogenesis^74, 75^. Cdc42 can be activated by multiple GEFs, which define the timing and location of its activity^76^. After being activated, Cdc42 executes its function through a series of effectors. We show here that in the context of branched actin assembly in neuronal growth cones, Asef1 is a key upstream Cdc42 activator, while N-WASP is a key downstream Cdc42 effector. Previous studies also implicated Cdc42^75, 77, 78^ and N-WASP^79–81^ functions in neurite outgrowth. Our findings that either knockdown or pharmacological inhibition of Cdc42 results in smaller growth cones conflict with an increased growth cone size found in hippocampal neurons isolated from Cdc42 knockout mice^74^. This discrepancy could be explained by some compensation effect after chronic ablation of Cdc42 in mice.

It is somewhat surprising that N-WASP plays so profoundly important roles in the assembly of virtually all branched actin filaments in growth cones, because another NPF, the WAVE regulatory complex (WRC), acting downstream of Rac1, is classically responsible for the formation of lamellipodia in most cell types^15, 82^. N-WASP, on the other hand, is typically implicated in quite local actin branching events, such as at clathrin-coated pits^83^, at the core of podosomes^84^ and invadopodia^85^ or at the base of filopodia^86^. However, Cdc42 also acts as an upstream activator of Rac1 through its other effectors, such as the Par3 – Par6 – aPKC polarity complex^87^. Therefore, active Rac1 through WRC can amplify the effects of Cdc42 and N-WASP on branched actin formation leading to expansion of the initial small focus of Arp2/3 complex activity over a larger area. Indeed, previous studies have found that both Cdc42 and Rac1 contribute to neurite outgrowth^75, 88^.

## Conclusion

We show here that APC mediates branched actin formation in neuronal growth cones through the Asef1 – Cdc42 – N-WASP pathway and that this pathway is likely also functional in other cell types. The initial formation of branched actin networks triggered by APC at dynamic microtubule tips can be further amplified and stabilized though other signaling events. Besides the Cdc42 – Rac1 crosstalk mentioned above, the PAR polarity complex, which is recruited by active Cdc42, can promote further APC clustering, thus forming a positive feedback loop^89^. Another Cdc42 effector and an ARD binding partner, IQGAP1, can stabilize the active Cdc42 state^90^ to prolong this signaling. On the other hand, some negative feedback loops, such as through activation of actomyosin contractility via the Cdc42 – MRCK – myosin II pathway^91^, can limit protrusion along the originally chosen narrow path. Together, this intricate signaling hub centered on APC has a capability to drive persistent neurite outgrowth in a particular direction or define the migration trajectory of other cells.

By revealing the molecular mechanism by which APC initiates branched assembly in cellular context, our study answers the fundamental question of how microtubules instruct the direction of leading-edge protrusion for directional cell migration. Moreover, our findings may also influence the future developments of treatment strategies for diseases caused by mutations of either APC or its downstream effectors.

## MATERIALS AND METHODS

### Cell Culture

Dissociated rat embryo hippocampal neurons isolated as described previously^92^ were obtained from the Neurons R Us Cell Service Core (University of Pennsylvania, Philadelphia, PA). In brief, hippocampi were dissected from brains of Sprague-Dawley rat embryos at embryonic day 18-20 and dissociated into individual cells by incubating in a trypsin-containing solution. The cells were then washed and plated on poly-D-lysine-coated glass coverslips (#GG-12-PDL, Neuvitro) at a concentration of 120,000 cells per 35-mm dish in 1.5 ml neurobasal medium (#21103049,ThermoFisher Sci.) with 2% B27 supplement (#17504044, ThermoFisher Sci.).

REF52 rat embryo fibroblasts and HEK293T cells (#CRL-11268, ATCC) were cultured in high glucose DMEM (#10569010, ThermoFisher Sci.) supplemented with 10% FBS (#F2442, Sigma) at 37°C and 5% CO_2_. Cells were plated on glass coverslips for fixed cell experiments or on MatTek 35 mm glass-bottomed dishes (#P35G-1.5-14-C, MatTek Inc.) for live cell imaging. CO_2_-independent phenol-red-free L15 medium (#21083027, ThermoFisher Sci.) supplemented with 10% FBS was used for live cell imaging. All cell lines were regularly tested for mycoplasma with the Mycoplasma Detection Kit (InvivoGen, Cat#: rep-mys-50).

### Antibodies

The following primary antibodies were used: APC-M2 rabbit polyclonal antibody against amino acids 959-1338 of APC ^93^ (a gift from Dr. Kristi Neufeld) was used at 1/50 dilution for immunofluorescence staining; Asef1 rabbit polyclonal antibody (#55213-I-AP, Proteintech) was used at 1/100 dilution for immunofluorescence staining and at 1/1000 dilution for Western blotting; Cdc42 rabbit polyclonal antibody (#2462, Cell Signaling Technology) was used at 1/500 dilution for Western blotting; N-WASP rabbit monoclonal antibody (clone 30D10, #4848S, Cell Signaling Technology) was used at 1/100 for immunofluorescence staining and at 1/1000 dilution for Western blotting; α-tubulin mouse monoclonal antibody (clone DM1α, #T-6199, Sigma) was used at 1/5000 dilution for Western blotting; β3-tubulin mouse monoclonal antibody (clone TUJ1, # 801201, BioLegend) was used at 1/200 dilution for immunofluorescence staining; and Cdc42 mouse monoclonal antibody (#610928, BD Transduction Laboratories) was used at 1/100 dilution for immunofluorescence staining. Secondary antibodies (used at 1/200 dilution) and phalloidin (used at 1/200 dilution) fluorescently labeled with Alexa Fluor-488, -594 and -680 were from Invitrogen.

### Constructs

The following expression constructs were used: YFP-APC (human) was a gift from E. M. Wenzel and J. Behrens^94^, pLL-mCherry and pLL-EGFP lentiviral vectors (pLL5.0) and pLL-EGFP-EB3 (human) construct were gifts from A. Efimov (Fox Chase Cancer Center, Philadelphia, PA)^95^. Full length pLL-EGFP-APC and its deletion mutants were generated by PCR from the YFP-APC template and cloned into pLL-EGFP-5.0 vector. N-APC and C-APC used in REF52 rescue experiment were cloned into C2-GFP vector (Clontech). The primers for full length APC (amino acids 1-2843) were 5’-TTCGTGACCGCGGCCAGATCTG-3’ (forward with SacII restriction site) and 5’-CAAGGGATCCACAGATGTCACAAGGTAAGACCC-3’ (reverse with BamHI restriction site). For N-APC (amino acids 1-958), the primers were 5’-TTCGTGACCGCGGCCAGATCTG-3’ (forward with SacII restriction site) and 5’-GAATGGGCCCCAATCGAGGGTTTCATTTGAC-3’ (reverse with ApaI restriction site). For C-APC (amino acids 2167-2843) the primers were 5’-GTAATCCGCGGCCAATGATTCTAAAACCAGGG-3’ (forward with SacII restriction site) and 5’-CAAGGGATCCACAGATGTCACAAGGTAAGACCC-3’ (reverse with BamHI restriction site). Primers to delete amino acid residues 959-2166 to produce mini-APC were designed with QuikChange Primer Design from Agilent. The forward primer was 5’-TATGCCAAATTAGAATACAAGAGAATTCTAAAACCAGGGGAGAAAAGT-3’ and the reverse primer was 5’-ACTTTTCTCCCCTGGTTTTAGAATTCTCTTGTATTCTAATTTGGCATA-3’. The PCR product was treated with DpnI enzyme and transformed into DH5α cells. F-tractin–mCherry and F-tractin–EGFP in lentiviral expression vectors were described previously^36^.

The shRNA sequence to target rat APC (sh#1) was described and validated previously, as well as the Luciferase-targeting sequence that was used as non-targeting control^36^. The other shRNA sequences targeting rat proteins were the following: 5’-AAGCAGACTTCCAGATCTATTCGGAGTACTG - 3’ for Asef1; 5’-CCTGATATCCTACACAACAAA - 3’ for Cdc42; and 5’- AAGACGAGATGCTCCAAATGG - 3’ for N-WASP. Oligos for shRNA construction were obtained from Integrated DNA Technologies, annealed and cloned into HpaI and XhoI restriction enzyme sites of the lentiviral pLL-GFP 3.7 vector (a gift from Dr. Luk Parijs, #11795, Addgene).

### Knockdown and rescue experiments with REF52 cells

REF52 cells were replated ∼16h before infection with shAPC/Cerulean-encoding lentivirus mixed with 5 µg/ml Protamine Sulfate (#P4020, Sigma). Culture medium was replaced after 16-18h of incubation of cells with the virus. REF52 cells were cultured for additional 2 days, trypsinized, collected by centrifugation at 1,200 rpm for 5 min and electroporated with the APC constructs in the C2 vector using Neon TM transfection system (#MPK5000, Invitrogen) using the following parameters: voltage 1350V, width 30ms, pulses 1. Electroporated cells were cultured for additional 2 days on glass coverslips before fixation.

### Lentivirus production and neuron infection

For producing lentiviral particles, expression plasmids containing shRNA sequences were co-transfected together with helper plasmids MD2G (a gift from Dr. Didier Trono, #12259, Addgene) and DVPR (a gift from Dr. Bob Weinberg, #12259, Addgene) into HEK-293T cells using Lipofectamine 3000 (#L3000008, Invitrogen). Virus was harvested on the 3rd day after transfection, filtered through 0.45 µm filter and applied to hippocampal neurons together with 5 µg/ml Protamine Sulfate (#P4020, Sigma) at ∼20 hrs after plating. Culture medium was replaced after 6-8 h of incubation of neurons with the virus. Neurons were cultured for additional 4 days before fixation.

### Light microscopy

Immunofluorescence staining was performed after fixation with 4% formaldehyde (Electron Microscopy Sciences) in PBS for 15 min followed by permeabilization with 0.1% Triton in PBS for 10 min. Confocal fluorescence microscopy was performed using Nikon Eclipse Ti-E inverted microscope (Nikon) equipped with CSU-X1 spinning disk (Yokogawa), dry PlanApo 20 × 0.75 NA, oil CFI60 Plan Apochromat Lambda 60x × 1.4 NA, and oil CFI60 Apochromat TIRF 100 × 1.49 NA objectives and QuantEM 512SC digital camera (Photometrics) and ORCA-FLASH 4.0 LTS+ sCMOS cameras. The system was driven by NIS-Elements software (Nikon).

### Western Blotting

Cells were lysed using cold lysis buffer containing 20 mM Tris-HCL (pH 7.5), 150 mM NaCl, 1 mM sodium EDTA, 1 mM EGTA, 1% NP-40, 1% sodium deoxycholate, 2.5 mM sodium pyrophosphate, 1 mM β-glycerophosphate, 1 mM Na_3_VO_4_, 1 μg/ml leupeptin, 1 mM PMSF and protease inhibitor cocktail (#P2714-1BTL, Sigma). Following brief sonication, lysates were clarified by centrifugation at 16,000g for 10 min at +4°C, protein concentration in supernatants was determined using Pierce^TM^ Bradford Assay Kit (#23246, Thermo Scientific). Protein samples were mixed with Laemelli Sample buffer (#1610747, BIO-RAD) and 5% 2-Mercaptoethanol and heated at 90°C for 10 min. About 20 µg (for cell lysates) of total protein was loaded onto NuPAGE 4-12% BT gradient gels (#NP0335, Invitrogen), resolved by SDS-PAGE, and transferred to 0.45 µm nitrocellulose membrane in NuPAGE Transfer Buffer for 90 min at 20V using the XCell SureLock Mini-Cell Electrophoresis and Western Blotting System (Invitrogen). The membrane was blocked with 5% dry milk in Tris buffer saline with Tween20 (TBST) (#9997S, Cell Signaling Technology) for 1 h before incubating with primary antibodies in the same blocking solution overnight at 4°C. The blots were probed with horseradish peroxidase-conjugated secondary antibodies at room temperature for 1h, developed with ECL Western Blotting Substrates (#RPN2109, Amersham; #R1002, Kindle Biosciences) and imaged using KuikQuant Imager (Kindle Biosciences) and ChemiDoc MP Imaging System (Bio-Rad).

### Platinum Replica Electron microscopy (PREM)

Sample preparation for PREM was performed as described previously^96, 97^ with minor modifications. In brief, cultured neurons were detergent-extracted and then fixed sequentially with 2% glutaraldehyde in 0.1 M Na-cacodylate buffer (pH 7.3), aqueous 0.1% tannic acid, and aqueous 2% uranyl acetate; critical point dried; coated with platinum and carbon; and transferred onto electron microscopic grids for observation. Detergent extraction was done with 0.5% Triton X-100 in PEM buffer (100 mM PIPES-KOH, pH 6.9, 1 mM MgCl_2_, and 1 mM EGTA) containing 4 μM phalloidin, and 10 μM taxol for 3 min at room temperature. Samples were analyzed using JEM 1011 transmission electron microscope (JEOL USA) operated at 100 kV. Images were captured with an ORIUS 832.10W CCD camera (Gatan) and presented in inverted contrast. Color labeling was performed using Hue/Saturation Adjustment Layer tool in Adobe Photoshop.

### Drug treatment

Cdc42 inhibitor ML141 (#M425250, Toronto Research Chemicals) and N-WASP inhibitor wiskostatin (#ab141085, Abcam) were dissolved in dimethyl sulfoxide (DMSO) to a stock concentration of 20 mM and 6 mM (1000X), respectively, diluted in culture medium to a final concentration of 20 μM (ML141) and 6 μM (wiskostatin), and added to primary neurons for 18-20 hrs after plating neurons for 4-6 hours. Neurons were then fixed for light or electron microscopy as described above.

### Image Analysis and Statistics

Image analyses and measurements were done using Fiji (ImageJ, NIH) or Adobe Photoshop software. GraphPad Prism version 8.1.1 for Windows was used for statistical analyses; graphs were generated using GraphPad or Excel.

For determination of the circularity index of REF52 cells transfected with EGFP-tagged APC constructs, cells were fixed 48 h after transfection and stained with GFP antibody to amplify the signal. Expressing cells were thresholded based on the GFP channel and circularity of the regions of interest was determined using Fiji.

Quantification of protrusions induced by APC-positive microtubules was described previously^36^. Briefly, kymographs from time-lapse sequences of REF52 cells coexpressing N-APC or mini-APC and F-tractin in different colors were generated along a straight or segmented line drawn along the path of a moving APC signal using Fiji. Cell protrusion events were scored positive if they began simultaneously with or within 30 sec after the APC track touched the cell edge or the actin-rich zone. Only tracks that reached the cell edge were taken into consideration. The quantifications for full-length APC and EB3 in figure 1M are reproduced from our previous study for comparison^36^.

For quantification of phalloidin fluorescence intensity in growth cones and of growth cone area, neurons were imaged by spinning disk confocal microscopy at 60x magnification. Analysis was performed using Fiji software after summing up confocal slices, background subtraction and thresholding of individual growth cones.

Neurite length quantification was performed using immunofluorescence images of β3-tubulin antibody staining acquired at 20x magnification. After background subtraction, neurite tracing and length measurement were performed using Fiji software. Entangled neurites were not included in the analysis.

Quantification of actin cytoskeleton architecture at neurite tips was performed using PREM images. Every neurite tip without technical defects or extensive overlap with other structures was included in the analysis. For determination of the presence of branched actin network at the neurite tip, each neurite tip was traced back for 20 μm to determine the presence of branched actin patches. Branched actin networks were recognized by a combination of some or all of the following characteristics: the presence of frequent Y-shaped configurations, short filament lengths, a wide range of filament orientations, and abundance of actin filament ends within a local area of the actin network. Around 10 individual neurons were chosen in each treatment group, and all neurite tips from these neurons were pooled together for each treatment type for percentage calculation.

## ACKNOWLEDGMENTS

We thank Drs. C. Yang and A. Chougule for valuable discussions. We thank Drs. E.M. Wenzel, J. Behrens, S. Kojima, G.G. Borisy, A. Efimov, K. Wang, Z. Yu and Z. Qin for generous gifts of reagents and/or sharing their equipment. We thank Dr. M. S. Shutova for constructing the control luciferase-targeting shRNA. This work was supported by NIH grant R35 GM 140832 to T.M.S.

## AUTHOR CONTRIBUTIONS

X.F. and N.E. performed experiments and analyzed data; X.F. wrote the manuscript; T.M.S. oversaw research and edited the manuscript.

## COMPETING INTERESTS

Authors declare no competing interests.

---

**Figure. S1.**
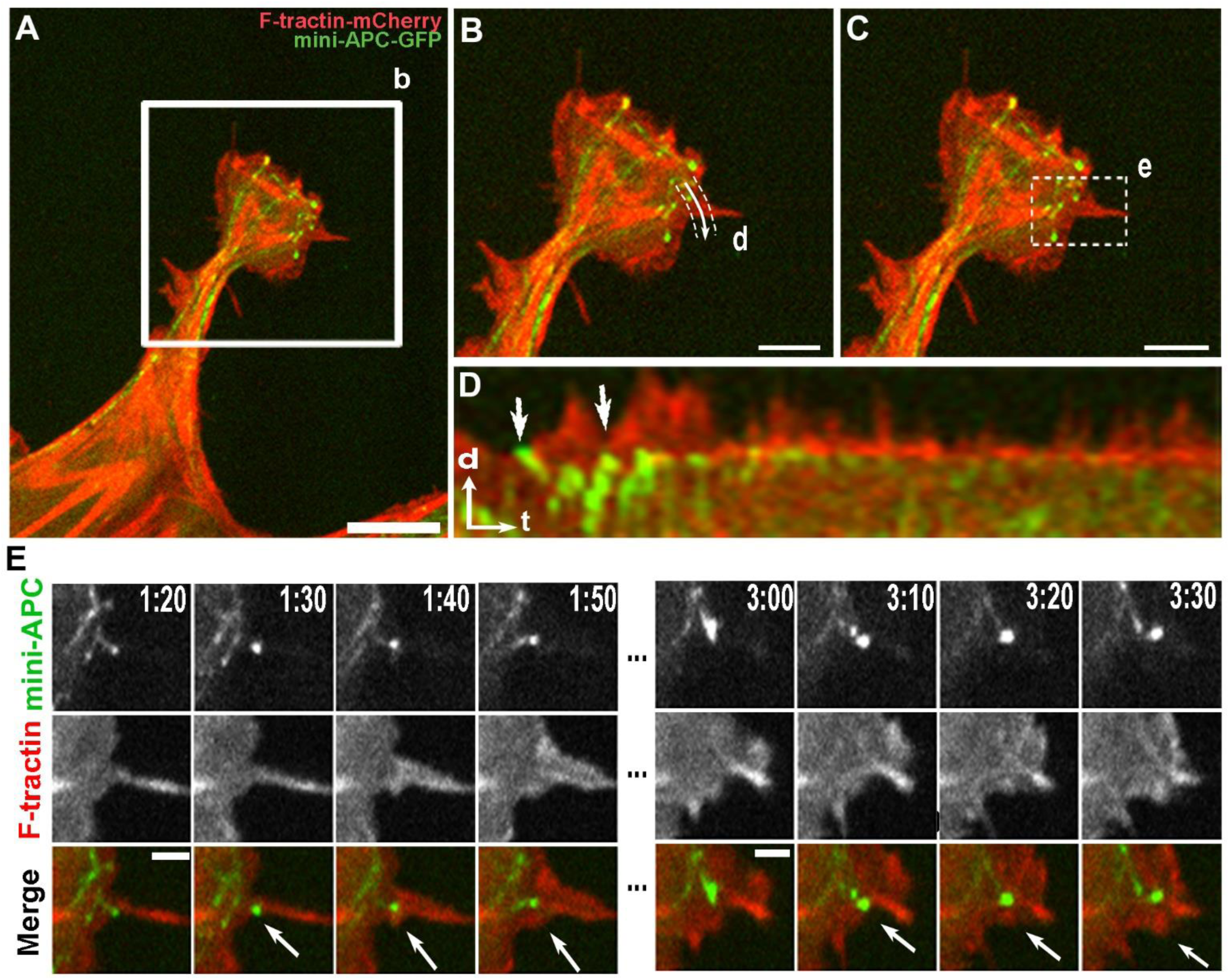
Dynamics of mini-APC in REF52 cells. (A) Process of REF52 cell expressing GFP-mini-APC (green) and F-tractin-mCherry (red). Boxed region is enlarged in B. (B) Enlarged view of the process tip showing the linear path (d) for kymograph shown in D. (C) Same enlarged view of the process tip as in B showing the region of interest (e) for time-lapse frames in E. (D) Kymograph of mini-APC-GFP (green) and F-tractin-mCherry (red) along the path shown in panel B. Encounters of the mini-APC punctum with the cell edge (arrows) are followed by F-actin protrusions. (E) Time-lapse frames for the boxed region in panel C. Arrows mark the actin-rich protrusion formed after the mini-APC-GFP encounters the cell edge. Time is shown in min:sec. Scale bars: 10 µm (A); 5 µm (B, C), and 2 µm (E). Arrows in D: time (t) = 60 s; distance (d) = 2 µm.

**Figure. S2.**
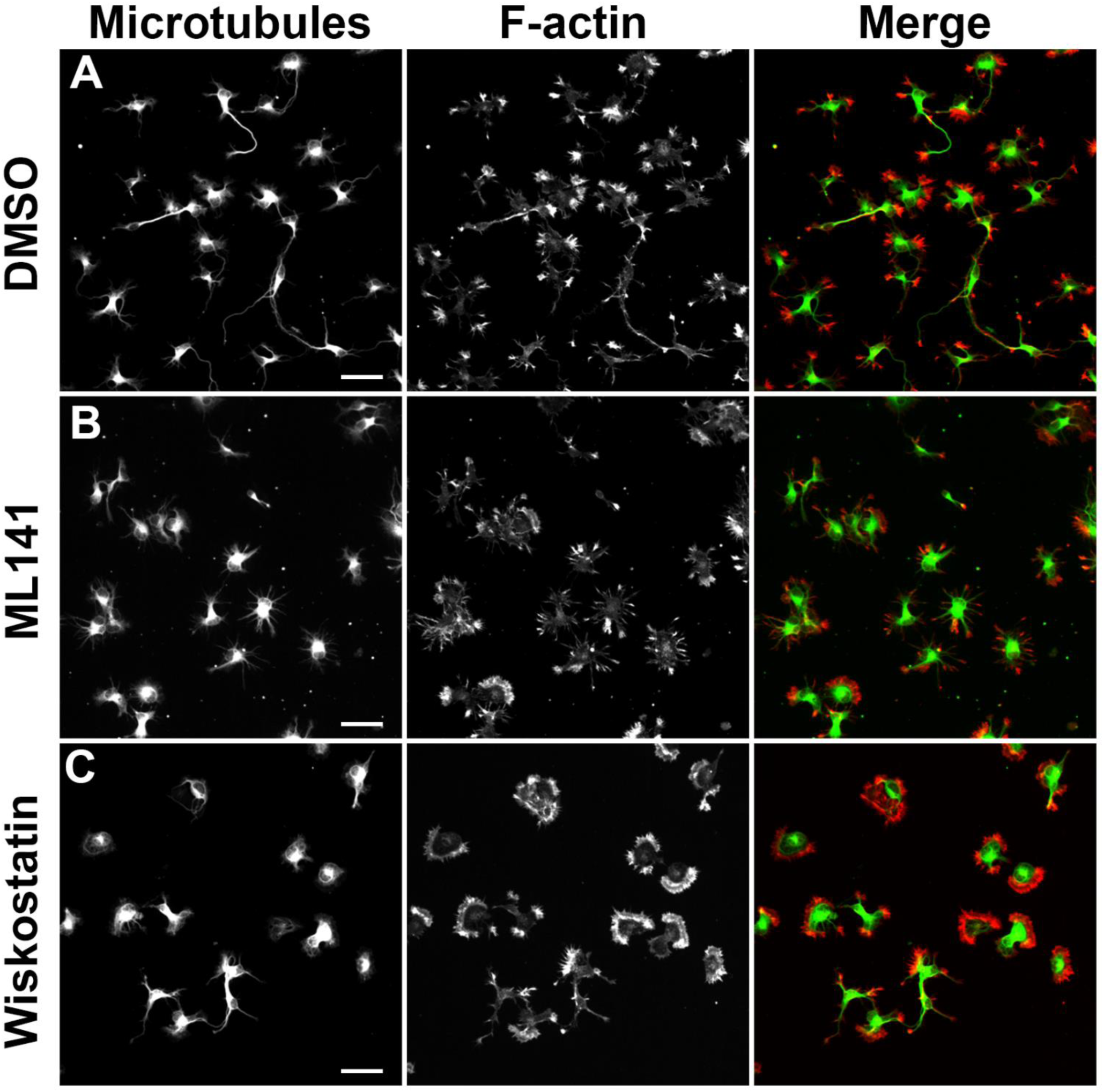
Pharmacological inhibition of Cdc42 and N-WASP reduces neurite lengths in young rat hippocampal neurons. Control DMSO-treated neurons (A), ML141-treated neurons (DIV1) (B) and wiskostatin-treated neurons (C) were stained with β3-tubulin antibody (TUJ1, green) and phalloidin (red). Scale bars: 50 µm.

**Video. 1.**
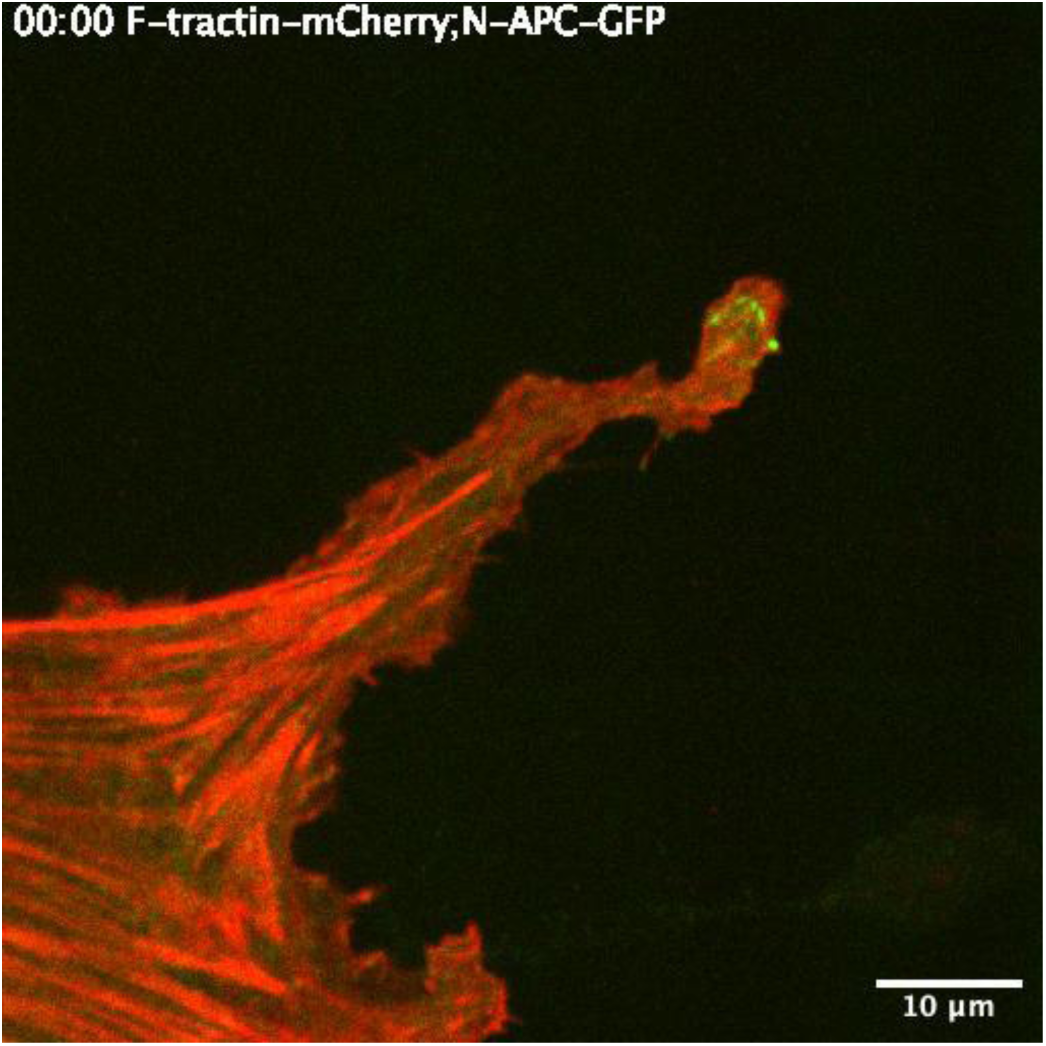
Dynamics of GFP-N-APC (green) and mCherry-F-tractin (red) in REF52 cells. The total length of the sequence is 8 min 30 s with 10.29 sec intervals between frames. Scale bar, 10 µm.

**Video. 2.**
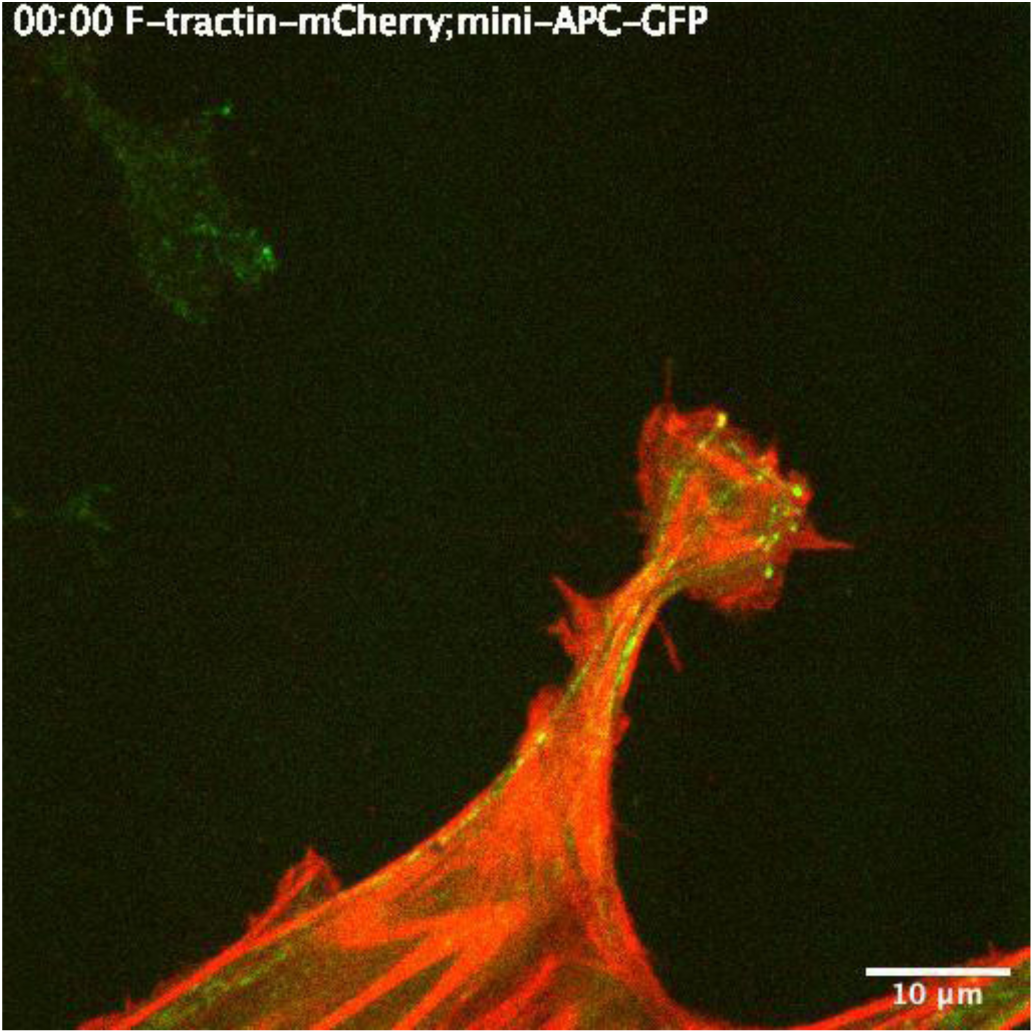
Dynamics of GFP-mini-APC (green) and mCherry-F-tractin (red) in REF52 cells. The total length of the sequence is 8 min 20 s with 10.29 sec intervals between frames. Scale bar, 10 µm.

## Notes

### Competing Interest Statement

The authors have declared no competing interest.

